# Prediabetes Phenotype Clusters in the Diabetes Prevention Program Study

**DOI:** 10.1101/2024.12.02.626435

**Authors:** Benjamin M. Stroebel, Meghana Gadgil, Kimberly Lewis, Kayla Longoria, Li Zhang, Elena Flowers

## Abstract

**Objective:** The purpose of this study was to apply clustering methods to identify and characterize prediabetes phenotypes and their relationships with treatment arm and type 2 diabetes (T2D) outcomes in the Diabetes Prevention Program (DPP), and to compare the utility of additional clustering measures in phenotype characterization and T2D risk stratification.

**Research Design and Methods:** This was a secondary analysis of data from a subset of participants (n=994) from the previously completed Diabetes Prevention Program trial. Unsupervised k-means clustering analysis was applied to derive the optimal number of clusters of participants based on common clinical risk factors alone or common risk factors plus more comprehensive measures of glucose tolerance and body composition.

**Results:** Five clusters were derived from common clinical characteristics and the addition of comprehensive measures of glucose tolerance and body composition. Within each modeling approach, participants show significantly different levels of risk factors. The clinical only model showed higher accuracy for time to T2D, however the more comprehensive models further differentiated a metabolically health overweight phenotype. For both models, the greatest differentiation in determining time to T2D was in the metformin arm of the trial.

**Conclusions:** Data driven clustering of patients with prediabetes allows for identification of prediabetes phenotypes at greater risk for disease progression and responses to risk reduction interventions. Further investigation into phenotypic differences in treatment response could enable better personalization of prediabetes and T2D prevention and treatment choices.

## INTRODUCTION

The global prevalence of type 2 diabetes (T2D), prediabetes, and related comorbidities continues to increase. (1) Currently, determination of prediabetes, which is a major risk factor for T2D, relies on the use of either hemoglobin A1c (HbA1c) or fasting blood glucose (FBG), (2) assigning patients a binary classifier. However, significant mechanistic heterogeneity has been demonstrated in both T2D and prediabetes, with recent studies identifying sub-phenotypes that may confer different degrees of risk for T2D progression and adverse outcomes. (3-10) This evidence suggests that the binary classification approach is inadequate for detecting potential differences in the underlying etiology, progression to T2D, variability in disease severity and progression, and the potential for responses to risk reduction interventions. Advancements in the characterization and understanding of the heterogeneity of T2D and prediabetes have the potential to increase precision in risk assessment and treatment.

Several prior studies identified sub-phenotypes of individuals with T2D based on clustering of clinical variables, (3-8; 11) which may have implications for treatment, management, and prevention of diabetes-related complications. However, few prior studies focused on cluster phenotypes in the prediabetes window, during which time intensive risk reduction interventions have the potential to prevent T2D and related complications. In cohorts from the United Kingdom and Sweden, individuals with prediabetes were clustered according to 2-hour plasma glucose, insulin secretion, fasting insulin, waist circumference, hip circumference, BMI, HDL cholesterol (HDL-C), and triglycerides. These studies resulted in the identification of six phenotypes among individuals with prediabetes, with differing risk for development of T2D and chronic kidney disease (CKD). (6) This approach was repeated in a more diverse cohort from the National Health and Nutrition Examination Survey (NHANES) study. Individuals with prediabetes were clustered according to age at prediabetes diagnosis, BMI, HbA1c, FBG, 2-hour plasma glucose, HOMA-IR, HOMA2-B, triglycerides, HDL-C, aspartate transaminase (AST), alanine transaminase (ALT), and glutamyl-transpeptidase (GGT), resulting in six clusters with differing cardiac and renal outcome risk. (7)

While all of the prior studies included common measures of body composition, none included more granular estimates of adipose tissue depots. Visceral adipose tissue is known to be metabolically more harmful than subcutaneous adipose tissue, (12; 13) but differentiation between the two is not readily measured by BMI or waist circumference (14). Addition of measures of ectopic fat depot mass has the potential to further differentiate sub-phenotypes of individuals with prediabetes and associated risk for progression to T2D. There is the possibility for a proxy measure for the underlying body composition risk profile that can be identified using specific combinations of common risk factors based on underlying statistical associations with these more granular body composition measures.

Prior studies have also not evaluated whether sub-phenotypes are related to responses to risk reduction interventions. It is increasingly recognized that there are differences in individual responses to interventions that result from both biological (e.g., genetic) and social-behavioral (e.g., food insecurity) characteristics. These more granular sub-phenotypes have the potential to elucidate not only risk profiles that result from individual differences but also likelihood of response to a given risk reduction intervention.

The Diabetes Prevention Program was a landmark clinical trial that compared an intensive lifestyle intervention and metformin to placebo for prevention of incident T2D. (15) Building on emerging evidence about sub-phenotypes, the present study will apply clustering methods to identify and characterize prediabetes phenotypes that include measures of ectopic fat. We also evaluated relationships between these clusters and treatment arm and incident T2D in the DPP trial.

## RESEARCH DESIGN AND METHODS

### Participants and Study Design

This study was a secondary analysis of data from participants in the DPP trial, which has been described in detail. (16-18) Briefly, participants were recruited from 27 centers across the U.S. between 1996-1999, with oversampling from racially minoritized groups. (18) Inclusion criteria required participants to be >25 years of age, have a BMI >24 24 kg/m^2^, FBG 95-125 mg/dl, and a 2-hour post-challenge glucose 140-199 mg/dl. Exclusion criteria included use of medications known to alter glucose tolerance or serious illness. A total of 3,234 participants were enrolled. The study described in this manuscript includes a subset of participants (n=994) who underwent abdominal computerized tomography (CT) imaging and had measures of body composition available. We included data collected at baseline and diabetes outcome data collected throughout the DPP trial through the primary endpoint at two years.

### Demographic and Clinical Data Collection

DPP participant demographic characteristics, medical history, and FBG were collected by trained study personnel at the first screening visit. (18) Physical measurements, oral glucose tolerance tests, behavioral data, and other laboratory tests were collected at the second interview screening visit. (18) History and physical exams were collected at the third screening visit. (18) Blood was collected by venipuncture according to a standardized protocol.

### Statistical Analyses

All statistical analyses were performed using R (version 4.3.2). (19) Data were summarized using descriptive statistics (means/medians and standard deviations/interquartile ranges for continuous variables and counts and percentages for categorical variables). To compare between clusters, we used Chi-squared (categorical variables) or analysis of variance (ANOVA) (continuous variables) to evaluate the demographic and clinical characteristics of the participants. Box and whisker plots were created to visualize the relative differences in clinical variables by cluster.

Clustering model variables were selected based on *a priori* knowledge about T2D risk factors, variables included in other clustering analysis studies that were also available in the DPP sample, and mechanistic relationships to metabolism and insulin signaling. (3-8) We developed two models for clustering of participants. The clinical model included routinely measured clinical characteristics and risk factors (i.e., age, BMI, FBG, HbA1c, HDL-C, triglycerides, and waist circumference). The clinicalPLUS+ model included all the aforesaid variables, with the addition of measures for glucose tolerance and body composition (i.e., 2-hour glucose, alanine transaminase (ALT), aspartate transaminase (AST), HOMA-B, HOMA-IR, L2-L3 subcutaneous fat area, and L2-L3 visceral fat area) given their associations with mechanisms involved in body composition, metabolism, and insulin signaling. (3-8) All variables used in clustering models were collected at trial enrollment and were centered to a mean value of 0 and a standard deviation of one.

Unsupervised k-means clustering was conducted for both the clinical and clinicalPLUS+ variables for a range of cluster counts (k=1-10). The average silhouette width, which ranges from -1 to 1 (1 indicating well-matched clusters), was calculated for each K value using the factoextra package in R to assess cluster stability and similarity within each cluster. (20) The silhouette widths supported K values of two, four, and five (average silhouette widths 0.19, 0.18, and 0.16) in the clinical model, and two, three, and five (average silhouette widths 0.17, 0.13, 0.12) in the clinicalPLUS+ model as best fitting the data. We selected cluster counts of five for both the clinical and clinicalPLUS+ models, as this choice provided clearer delineation between cluster profiles across the clustering characteristics.

Five-cluster k-means models were subsequently created for both the clinical and clinicalPLUS+ models using the Cluster package in R. (21) Cluster labels were assigned based on mean values of each variable included in the clusters.

Kaplan-Meier (KM) survival estimates and overall log-rank tests with post-hoc tests adjusted for multiple comparisons via Bonferroni-Holm corrections were used to assess differences in time to T2D development between clusters in the clinical and clinicalPLUS+ models, both overall and by DPP treatment arm. Cox Proportional Hazard (CPH) models adjusted for DPP treatment, participant sex, race and ethnicity were used to determine hazard ratios (HR) for time to T2D at the primary trial endpoint of 2-years by cluster. K-fold cross-validation was used to compute c-index scores for the fully adjusted clinical and clinicalPLUS+ Cox models using the glmnet package for R. (22; 23)

## RESULTS

### Cluster Characteristics and Labels

Five distinct clusters were identified and labelled in the clinical cluster models: 1) <u>older protected</u>, with the highest proportion of females, highest HDL-C and second highest age, lowest BMI, triglycerides, and FBG; 2) <u>dyslipidemia</u>, the smallest cluster with the lowest proportion of females, characterized by the highest triglycerides and lowest HDL-C; 3) <u>insulin resistant</u>, with the highest age, FBG and HbA1c; 4) <u>younger protected</u>, the largest cluster, featuring the lowest age, FBG, and HbA1c, relatively lower BMI and waist circumference; and 5) <u>higher adiposity protected</u>, with the second highest proportion of females, highest BMI and waist circumference and relatively lower triglycerides (**Figure 1**). All clinical clusters differed significantly in their sex and race and ethnicity composition (**Supplementary Table 1**).

**Figure 1.**
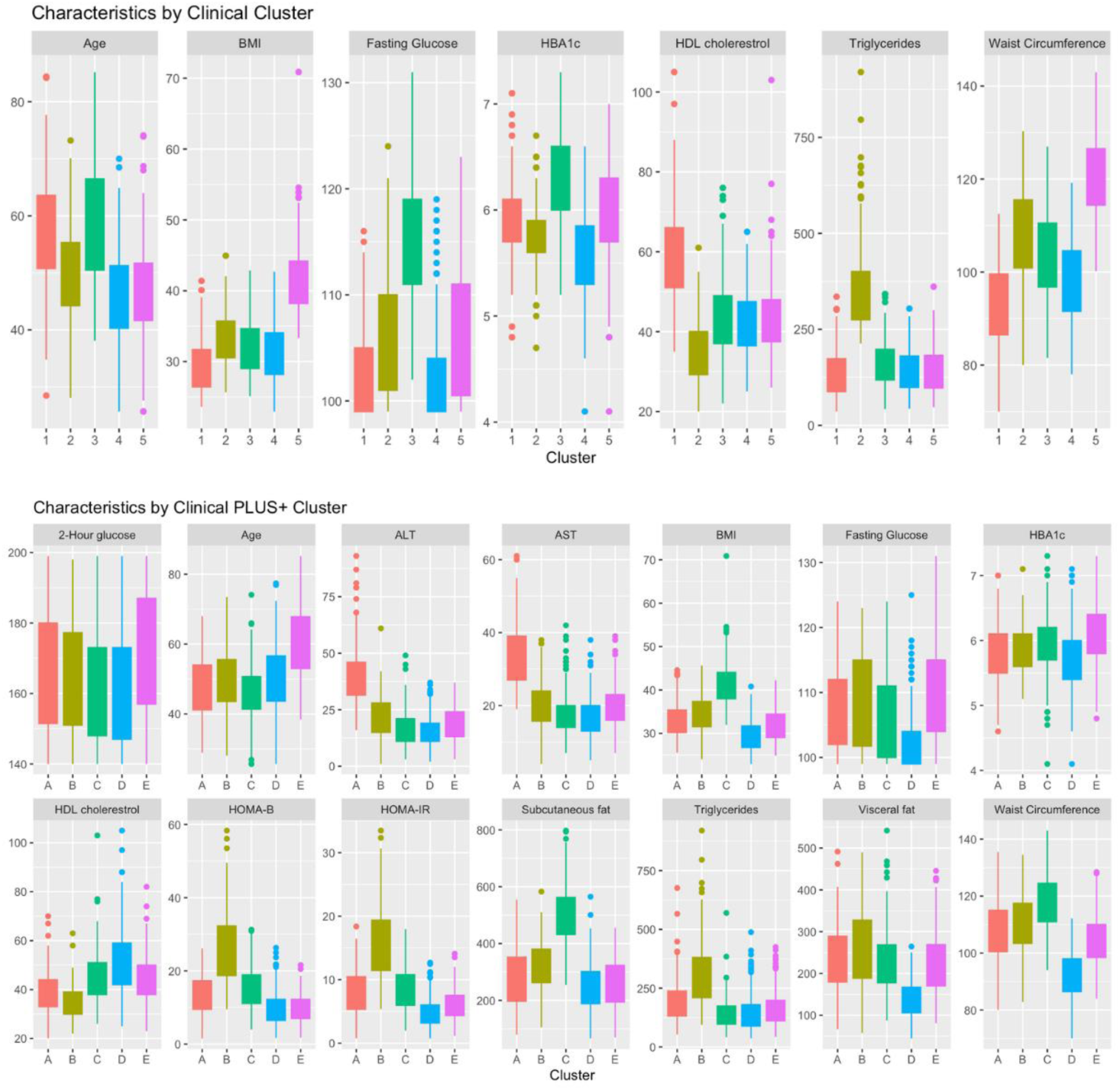
Clinical Characteristics by Cluster. Distribution of clinical risk factors included in phenotype models **Box and whisker plots:** Each plot shows the distribution of a clinical risk factor by phenotype cluster. Within each box, the bottom border represents the 25^th^ percentile and the upper border represents the 75^th^ percentile. The lowest point on the vertical line represents the minimum and the highest point represents the maximum. Small circles represent outliers. **Phenotype groups for the clinical model:** 1: older protected 2: dyslipidemia 3: insulin resistant 4: younger protected 5: higher adiposity protected **Phenotype groups for the clinicalPLUS+ model:** A: hepatic steatosis, B: dyslipidemia-insulin resistance C: subcutaneous adipose protected D: protected E: older dysglycemia **Y-axis shows units as described below:** 2- hour glucose - mg/dL age - years alanine transaminase (ALT) - IU/L aspartate aminotransferase (AST) - U/L body mass index (BMI) - kg/m^2^ for BMI fasting glucose – mg/dL hemoglobin A1c (HbA1c) - % high density lipoprotein (HDL) cholesterol – mg/dL homeostasis model assessment B (HOMA-B) homeostasis model assessment insulin resistance (HOMA-IR) subcutaneous fat – cm^2^/m^2^ triglycerides – mg/dL visceral fat – cm^2^/m^2^ waist circumference – cm

Five distinct clusters were also identified and labeled in the clinicalPLUS+ clustering models (labeled with letters rather than numbers to differentiate them from the clinical clusters): A) <u>hepatic steatosis</u>, the lowest proportion of females, lower HDL-C, higher triglycerides, high visceral fat area, and the highest AST and ALT; B) <u>dyslipidemia-insulin resistance</u>, the smallest cluster with the lowest HDL-C, and highest triglycerides, visceral fat area, HOMA-IR, and HOMA-B; C) <u>subcutaneous adipose protected</u>, a predominantly female cluster with the lowest age, highest BMI and subcutaneous fat area, lowest triglycerides and 2-hour glucose, with higher HOMA-measures and HbA1c; D) <u>protected</u>, the largest cluster with the highest proportion of women and the lowest BMI, waist circumference, visceral and subcutaneous fat areas, FBG, AST, ALT, HbA1c, HOMA-measures, and highest HDL-C; and E) <u>older dysglycemia</u>, the oldest cluster featuring lower BMI, waist circumference, HOMA-IR, and subcutaneous fat area, and the highest FBG, 2-hour glucose, HbA1c, and HOMA-B (**Figure 1**). All clinicalPLUS+ clusters differed significantly in their sex and race and ethnicity composition (**Supplementary Table 1**). The comparisons in cluster membership between clinical and clinicalPLUS+ are shown in **Supplementary Table 2**.

### Time to Incident T2D – Clinical Clusters

Time to T2D differed significantly between clinical clusters, both overall and by DPP treatment arm, In univariate KM analyses (**Figure 2**, p<0.05). Post-hoc tests indicated that time to T2D development was significantly lower overall in several comparisons: insulin resistant (cluster 3) vs older protected (cluster 1), older protected (cluster 1) vs younger protected (cluster 4), insulin resistant (cluster 3) vs dyslipidemia (cluster 2), dyslipidemia (cluster 2) vs younger protected (cluster 4), insulin resistant (cluster 3) vs younger protected (cluster 4), insulin resistant (cluster 3) vs higher adiposity (cluster 5), and higher adiposity protected (cluster 5) vs younger protected (cluster 4) (adjusted p<0.05). Time to T2D was also significantly lower in the placebo arm in the insulin resistant (cluster 3) vs younger protected (cluster 4) and in higher adiposity protected (cluster 5) vs younger protected (cluster 4) (adjusted p<0.05). In the metformin arm, time to T2D was significantly lower in insulin resistant (cluster 3) vs younger protected (cluster 4) (adjusted p<0.05). Time to T2D did not differ significantly between phenotype clusters in the lifestyle arm in post-hoc tests (adjusted p>0.05). Adjusted Cox models showed higher HR (95% CI) for T2D, relative to younger protected (cluster 4), in older protected (cluster 1), (2.22 (1.20, 4.11)), dyslipidemia (cluster 2) (2.32 (1.13, 4.77)), insulin resistant (cluster 3), (4.93 (2.77, 8.75)) and higher adiposity protected (cluster 5) (2.76 (1.51, 5.03)) (**Table 1**). The fully adjusted CPH model with clinical clusters achieved a Brier score of 0.100, a c-index score of 0.68, and the proportional hazards assumption was met. In terms of overall prevalence, the highest risk groups had 3-fold higher estimates compared to the lowest risk groups (e.g., 26% in the insulin resistant (3) group compared to 6% in the youngest and lowest risk group) (**Supplementary Table 3**).

**Figure 2.**
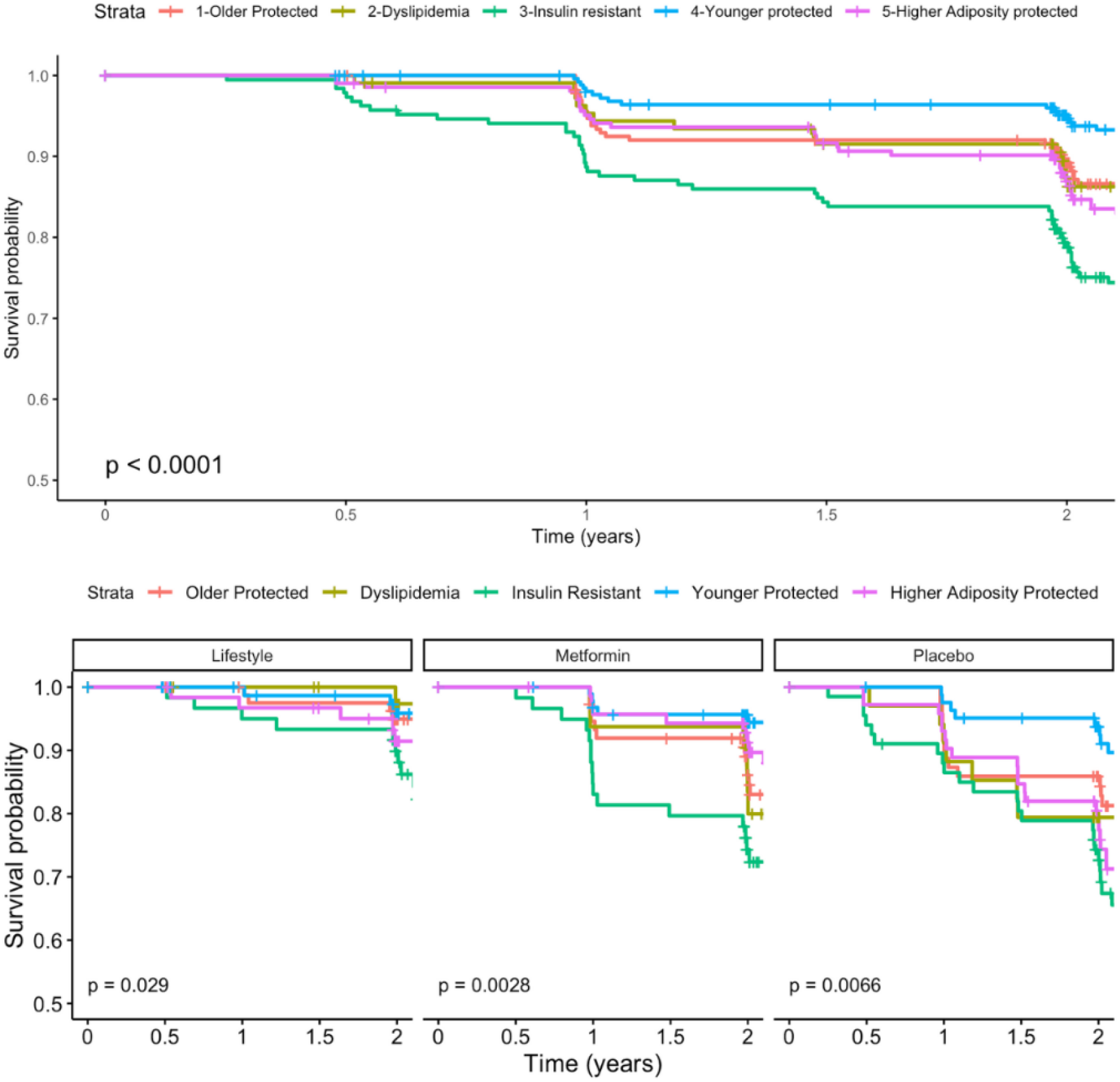
Time to Incident T2D – Clinical Clusters. Survival curves for incident type 2 diabetes in the clinical cluster model Curves represent the proportion of the sample free from T2D at a given time point. Data were censored at the primary trial endpoint of 2-years.

**Table 1.**

| **CLUSTER** | **HR** | **95% CI** | **p-value** |
| --- | --- | --- | --- |
| **Clinical model** |  |  |  |
| Older Protected | 2.22 | 1.20, 4.11 | 0.0113 |
| Dyslipidemia | 2.32 | 1.13, 4.77 | 0.0215 |
| Insulin Resistant | 4.93 | 2.77, 8.75 | <0.0001 |
| Younger Protected | ref | na | na |
| Higher Adiposity Protected | 2.76 | 1.51, 5.03 | 0.001 |
| **Clinical PLUS+ model** |  |  |  |
| Hepatic Steatosis | 2.01 | 1.07, 3.77 | 0.0302 |
| Dyslipidemia-Insulin resistance | 1.92 | 0.96, 3.86 | 0.0644 |
| Subcutaneous-Adipose Protected | 1.77 | 1.07, 2.92 | 0.0258 |
| Protected | ref | na | na |
| Older Dysglycemia | 3.03 | 1.89, 4.86 | <0.0001 |

### Time to Incident T2D – ClinicalPLUS+ Clusters

Time to T2D differed significantly between clinicalPLUS+ clusters overall and in the lifestyle treatment arm (**Figure 3**, p<0.05). Post-hoc tests indicated that time to T2D development was significantly lower in older dysglycemia (cluster E) compared to protected (cluster D) (adjusted p<0.05). Adjusted Cox models indicate that the HR (95%CI) for T2D, relative to protected (cluster D), is 2.01 (1.07, 3.77) for hepatic steatosis (cluster A), 1.92 (0.96, 3.86) for dyslipidemia-insulin resistance (cluster B), 1.77 (1.07, 2.92) for subcutaneous adipose protected (cluster C), and 3.03 (1.89, 4.86) for older dysglycemia (cluster E) (**Table 1**). The fully adjusted CPH model with clinicalPLUS+ clusters achieved a Brier score of 0.123, a c-index of 0.61, and the proportional hazards assumption was met. As with the clinical models, an evaluation of total prevalence after 2-years showed drastically higher estimates in the higher risk groups (e.g., 21% in the older dysglycemia (E) cluster compared to 9% in the protected (D) cluster) (**Supplementary Table 3**).

**Figure 3.**
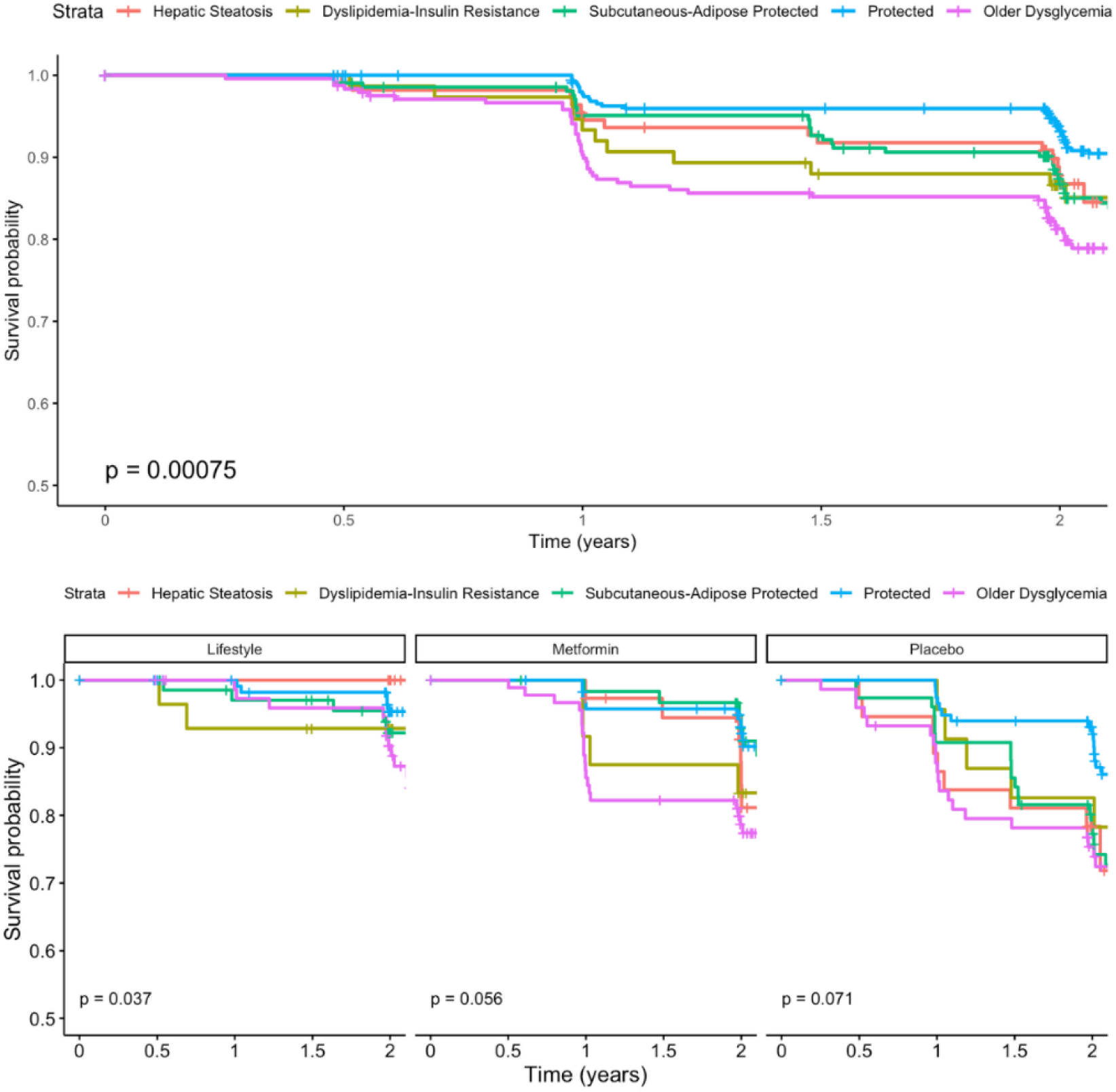
Time to incident T2D – Clinical PLUS+ Clusters. Survival curves for incident type 2 diabetes in the clinicalPLUS+ cluster model Curves represent the proportion of the sample free from T2D at a given time point. Data were censored at the primary trial endpoint of 2-years.

## DISCUSSION

Using data from the DPP trial, we applied an unsupervised clustering algorithm and identified two sets of five clusters among participants with prediabetes based on seven (clinical) and 14 (clinical PLUS+) phenotypic variables. Our study demonstrated distinct differences in phenotypic characteristics and rates of progression to T2D between clusters. Relative time to T2D progression also differed according to DPP treatment arm, suggesting the effects of treatment on time to T2D progression may differ according to baseline sub-phenotype. In addition, overall prevalence of T2D varied significantly between clusters, with the highest risk groups exhibiting a 3-fold increase compared to the lowest risk groups (**Supplementary Table 3**). Previously, based on a binary prediabetes classification, these participants were considered to have similar risk profiles. our findings highlight the potential value of identifying distinct subgroups to better capture the heterogeneity in prediabetes risk, which would allow for increased precision in risk assessment and treatment.

The clinical cluster model that relies on widely available variables than the clinicalPLUS+ model demonstrated a slightly better Brier score. Consequently, the clinical cluster model may be more easily replicated in other datasets, highlighting its potential clinical utility as a risk stratification tool, especially in settings without access to advanced body composition measures.

As expected, the insulin resistant cluster 3 shows the greatest risk for incident T2D. The identified clinical dyslipidemia phenotype (cluster 2) shares a trend of high levels of triglycerides coupled with low HDL-C, also observed in patients with metabolic syndrome, has a moderate progression to T2D that validates this risk profile as being important even in the absence of obesity. (24) Similarly, the higher-adiposity protected phenotype (cluster 5), with the highest BMI and waist circumference and more moderate FBG and HbA1c, resembles the phenotype of patients with metabolically benign obesity, and may inform treatment decisions regarding weight management, insulin sensitivity, and lipid status.(25) Finally, the older protected also shows a more moderate rate of progression to T2D, which is consistent with prior data that show a diagnosis of prediabetes in older age (i.e., >65 years) has a lower rate of progression to T2D compared to a younger age at diagnosis. (26)

Characterization of clinicalPLUS+ cluster phenotypes revealed significant differences in underlying metabolic states that could inform further studies on the mechanistic underpinnings of prediabetes and progression to T2D. Phenotypic differences in transaminases, body composition, insulin resistance, and beta cell function have been previously described in phenotypic clusters of individuals with prediabetes. (6; 7) Our study noted significant differences in the relative area of visceral fat and subcutaneous fat between clusters in the clinicalPLUS+ clustering model. By traditional measures of body composition (i.e., BMI, waist circumference), cluster C was the most overweight/obese. However, upon more granular differentiation of adipose tissue depots, the subcutaneous adipose protected phenotype (cluster C) exhibited elevated L2-L3 subcutaneous fat and a comparatively longer time to T2D relative to the dyslipidemia-insulin resistance phenotype (cluster B) which also showed higher BMI and waist circumference compared to other groups but had elevated L2-L3 visceral fat area (vs subcutaneous). These observed differences suggest a dynamic role for body composition, consistent with prior evidence that links visceral fat volume with that of pancreatic fat, which has previously been associated with altered insulin secretion in individuals with prediabetes. (6)

In both the clinical and clinicalPLUS+ clusters, metformin is markedly more impactful in clusters that are characterized by cardiometabolic risk factors other than insulin resistance and dysglycemia. For the clinical models, the older protected (cluster 1), dyslipidemia (cluster 2), and higher adiposity protected (cluster 5) show the greatest reduction in incident T2D in the metformin group compared to placebo, with relatively little improvement in the insulin resistant group (cluster 3). For the clinicalPLUS+ models, it is the hepatic steatosis (cluster A) and subcutaneous adipose protected (cluster C) that show the greatest reduction in incident T2D, with the dyslipidemia-insulin resistance (cluster B) and older glycemia (cluster E) clusters showing relatively little change. The mechanisms underlying the impact of metformin are not fully understood, and recent evidence suggests that this agent may have benefits beyond diabetes prevention. (27; 28) The observation that metformin response varies by sub-phenotype suggests the potential for individualized treatment approaches to optimize risk reduction. In contrast, for both clustering approaches, all groups show benefit from the intensive lifestyle intervention, which supports continued efforts to identify strategies for pragmatic implementation of lifestyle and behavior change interventions that may include policy and social-level implications.

Similar to results in a previous clustering analysis of NHANES participants with prediabetes, (7) the distribution of the clusters identified in this study varied by race and ethnicity. Cluster 2 in the Jiang et. al. analysis had a higher proportion of individuals categorized as Black and the highest BMI, which is similar to clinical cluster 5 (higher adiposity protected) in our study. Evidence suggests that the common measures of body composition (e.g., BMI, waist circumference) may not provide equal information about risk between race and ethnic groups. (29) For example, clinicalPLUS+ cluster C (subcutaneous adipose protected) had high BMI, waist circumference, and subcutaneous adipose tissue, but relatively lower visceral adipose tissue and is somewhat protected from incident T2D compared to the hepatic steatosis cluster (cluster A) that had lower measures of BMI and waist circumference, but higher visceral adipose tissue and a smaller proportion of people categorized as Black. Clusters 1 and 2 from the Jiang et. al. paper were the youngest with the highest proportions categorized as Mexican-American or Other Hispanic with cluster 1 showing relatively healthier insulin and glucose metabolism, whereas cluster 2 had some of the poorest measures of insulin and glucose metabolism. In our study, the younger protected cluster (cluster 4) had the highest proportion of people categorized as Hispanic, similar to Jiang et. al.’s cluster 1. The second highest proportions were in the dyslipidemia (cluster 2) and insulin resistant (cluster 3) clusters, which may overlap with Jiang’s cluster 2 and suggest that there may be important differences in risk for T2D based on sub- phenotypes within race and ethnic group categorization. These observations highlight important potential contributions to risk for T2D that may result from both genetic differences determined by geographical ancestry as well as the social implications of race and ethnic group categorization.

### Limitations

Selection of variables to include in the clustering models reflected previous studies, but an optimal set of clustering characteristics and the ideal clustering models have yet to be established. Current clustering methods primarily upon phenotypic variables, which may be confounded, mediated, or otherwise influenced by socioenvironmental factors related to T2D progression. Cluster assignment is likely dynamic rather than static, highlighting the need for further studies to understand how phenotypes may evolve over time or change in response to risk reduction interventions and socioenvironmental factors.

As efforts to individualize treatment and management of prediabetes continue, working towards more precise stratification of risk for progression to T2D becomes increasingly important. Data driven clustering of patients with prediabetes works towards this end, allowing for identification of sub-phenotypes at greatest risk for disease progression and responses to risk reduction interventions. This secondary analysis of participants from the DPP trial who had prediabetes applied a data-driven approach to identify two models to cluster participants based on phenotypic characteristics. The clinical clusters leverage commonly measured risk factors for T2D and differentiated between subgroups based on age, insulin metabolism and dyslipidemia profiles, and anthropometric measures of body composition. In general, the youngest group with the healthiest risk factor profile had significantly lower rates of T2D while the most insulin resistant group had the highest, regardless of trial arm. The addition of more refined estimates of glucose and insulin metabolism and adipose tissue composition confirmed prior studies that showed type of adipose tissue, and not just BMI, differentiates risk for T2D. Overall, the intensive lifestyle intervention was effective at decreasing risk for T2D regardless of phenotype cluster, whereas some clusters showed markedly better response to metformin compared to others. We also observed differences that suggest further research is needed to determine how identified sub-phenotypes of prediabetes are related to ancestry vs race and ethnic group categorization and risk for T2D. Further investigation into phenotypic differences in treatment response could enable better personalization of prediabetes and T2D prevention and treatment choices. This is especially salient given the addition of newer therapeutics like GLP-1 receptor agonists where cost may factor into patient acceptability when compared to older therapeutics which may still offer phenotype-specific efficacy. (30)

## Author Contributions

BMS performed all data analysis and primarily drafted the manuscript; MG contributed to interpretation of the findings and clinical implications and reviewed and approved the final manuscript; KL contributed to interpretation of the findings and reviewed and approved the final manuscript; KDL contributed to interpretation of the findings and reviewed and approved the final manuscript; LZ contributed to the analysis plan and reviewed and approved the final manuscript; EF conceived of the overall study design and oversaw all aspects of study implementation and reviewed and approved the final manuscript.

## DATA AVAILABILITY

All datasets analyzed during the current study are publicly available through the NIDDK data repository.

