## Supplementary Table 1 for "Prediabetes Phenotype Clusters in the Diabetes Prevention Program Study"

|  |  | **Clinical Model** | | | | | **Clinical PLUS+ Model** | | | | |
| --- | --- | --- | --- | --- | --- | --- | --- | --- | --- | --- | --- |
|  | **Overall (n=994)** | **Older Protected (clinical) (n=232)** | **Dyslipidemia (clinical) (n=110)** | **Insulin Resistant (clinical) (n=186)** | **Younger Protected (clinical) (n=259)** | **Higher Adiposity Protected (clinical) (n=207)** | **Hepatic Steatosis (n=111)** | **Dyslipidemia-Insulin resistance (n=76)** | **Subcutaneous-Adipose Protected (n=209)** | **Protected (n=357)** | **Older Dysglycemia (n=241)** |
| **Age** | 52 (11) | 57 (9) | 50 (9) | 59 (10) | 46 (9) | 47 (8) | 48 (9) | 50 (10) | 47 (8) | 50 (9) | 61 (9) |
| **BMI** | 33.4 (6.1) | 29.2 (3.6) | 33.6 (3.7) | 32.0 (4.0) | 31.3 (3.9) | 41.8 (5.0) | 33.3 (4.2) | 34.7 (4.6) | 41.5 (5.1) | 29.5 (3.6) | 31.8 (3.7) |
| **Waist Circumference** | 104 (13) | 93 (9) | 107 (10) | 104 (9) | 98 (9) | 121 (9) | 107 (11) | 110 (12) | 118 (10) | 92 (8) | 104 (9) |
| **L2-L3 visceral fat** | 199 (85) | na | na | na | na | na | 240 (79) | 256 (102) | 229 (78) | 136 (42) | 229 (80) |
| **L2-L3 subcue fat** | 314 (134) | na | na | na | na | na | 287 (104) | 319 (96) | 501 (107) | 248 (80) | 259 (84) |
| **Triglycerides** | 168 (99) | 131 (58) | 367 (131) | 161 (62) | 46 (9) | 142 (58) | 200 (102) | 324 (174) | 140 (64) | 142 (72) | 164 (74) |
| **HDL-C** | 46 (12) | 59 (11) | 36 (8) | 43 (9) | 31.3 (3.9) | 44 (10) | 39 (9) | 35 (8) | 45 (11) | 51 (13) | 45 (9) |
| **Fasting glucose** | 106 (7) | 102 (4) | 106 (6) | 115 (6) | 98 (9) | 106 (7) | 107 ( 6) | 109 (8) | 106 (7) | 102 (4) | 110 (8) |
| **2-Hour glucose** | 164 (17) | na | na | na | na | na | 165 (17) | 165 (15) | 161 (15) | 161 (16) | 170 (17) |
| **HbA1c** | 5.9 (0.5) | 5.9 (0.4) | 5.8 (0.3) | 6.3 (0.5) | 5.6 (0.4) | 6.0 (0.5) | 5.8 (0.5) | 5.9 (0.4) | 6.0 (0.5) | 5.7 (0.4) | 6.1 (0.5) |
| **HOMA-IR** | 7.1 (4.2) | na | na | na | na | na | 7.9 (3.3) | 15.8 (6.3) | 8.5 (3.2) | 4.9 (2.2) | 6.0 (2.5) |
| **HOMA-B** | 12.6 (7.2) | na | na | na | na | na | 13.6 (5.5) | 26.6 (11.2) | 15.3 (5.9) | 9.8 (4.4) | 9.6 (3.7) |
| **AST** | 19.9 (7.8) | na | na | na | na | na | 33.4 (8.6) | 20.4 (7.3) | 17.7 (6.0) | 17.0 (5.2) | 19.7 (5.6) |
| **ALT** | 19.8 (11.6) | na | na | na | na | na | 41.2 (14.3) | 21.3 (10.5) | 17.0 (8.2) | 15.1 (6.6) | 18.7 (7.0) |
| **% Female** | 67 | 81 | 55 | 48 | 66 | 6.0 (0.5) | 32 | 52 | 84 | 85 | 44 |
| **Race and Ethnicity** |  |  |  |  |  |  |  |  |  |  |  |
| **White** | 575 (58) | 131 (56) | 79 (72) | 91 (49) | 154 (59) | 120 (58) | 67 (60) | 46 (61) | 110 (53) | 202 (57) | 150 962) |
| **Black** | 213 (21) | 54 (23) | 6 (5) | 58 (31) | 36 (14) | 59 (29) | 13 (12) | 8 (11) | 64 (31) | 73 (20) | 55 (23) |
| **Hispanic** | 163 (16) | 31 (13) | 19 (17) | 29 (16) | 60 (23) | 24 (12) | 21 (19) | 17 (22) | 34 (16) | 61 (17) | 30 (12) |
| **Other** | 43 (4) | 16 (7) | 6 (5) | 8 (4) | 9 (3) | 4 (2) | 10 (9) | 5 (7) | 1 (0) | 21 (6) | 6 (2) |
