## Supplementary Table 2 for "Prediabetes Phenotype Clusters in the Diabetes Prevention Program Study"

|  | **Clinical PLUS+ Cluster** | |  |  |  |
| --- | --- | --- | --- | --- | --- |
| **Clinical Cluster** | Hepatic Steatosis | Dyslipidemia-Insulin Resistance | Subcutaneous-Adipose Protected | Protected | Older Dysglycemia |
| Older protected | 7 | 3 | 7 | 170 | 45 |
| Dyslipidemia | 27 | 39 | 5 | 16 | 23 |
| Insulin Resistant | 19 | 15 | 11 | 7 | 134 |
| Younger Protected | 41 | 7 | 20 | 164 | 27 |
| Higher Adiposity protected | 17 | 12 | 166 | 0 | 12 |
