## Supplementary Table 3 for "Prediabetes Phenotype Clusters in the Diabetes Prevention Program Study"

|  | **2-year Follow-up** | | | **Full Follow-up (mean 2.4 years, max 4.1 years)** | | |
| --- | --- | --- | --- | --- | --- | --- |
| **Clinical Clusters** | **cases (n)** | **censored (n)** | **prevalence (%)** | **cases (n)** | **censored (n)** | **prevalence (%)** |
| 1-Older protected | 29 | 203 | 12.5 | 36 | 196 | 15.5 |
| 2-Dyslipidemia | 14 | 96 | 12.7 | 18 | 92 | 16.4 |
| 3-Insulin Resistant | 48 | 138 | 25.8 | 56 | 130 | 30.1 |
| 4-Younger Protected | 16 | 243 | 6.2 | 20 | 239 | 7.7 |
| 5-Higher Adiposity Protected | 33 | 174 | 15.9 | 42 | 165 | 20.3 |
| **Clinical PLUS+ Clusters** |  |  |  |  |  |  |
| A-Hepatic Steatosis | 16 | 96 | 14.3 | 20 | 91 | 18.0 |
| B-Dyslipidemia-Insulin Resistance | 11 | 65 | 14.5 | 15 | 61 | 19.7 |
| C-Subcutaneous-Adipose Protected | 31 | 178 | 14.8 | 39 | 170 | 18.7 |
| D- Protected | 31 | 326 | 8.7 | 38 | 319 | 10.6 |
| E- Older Dysglycemia | 51 | 190 | 21.2 | 60 | 181 | 24.9 |
